# StainStyleSampler: Clustering-based sampling of whole slide image appearances

**DOI:** 10.1101/2025.04.04.647194

**Authors:** Maya Maya Barbosa Silva, Sabine Leh, Hrafn Weishaupt

## Abstract

The appearance of whole slide biopsy images is greatly affected by various factors such as laboratory procedures or the choice of digital slide scanners. The resulting variations in image styles within and across batches of histological images represent one of the major obstacles to the development of generalizable machine learning algorithms. To overcome this challenge, a lot of research has focused on stain normalization and stain augmentation techniques. While such approaches provide effective strategies to reduce stain variation or increase stain invariance, respectively, they typically involve only limited modeling or sampling of the underlying stain style distribution. Tools for a streamlined sampling of different aspects of such a distribution, which would be crucial e.g. for explicitly evaluating machine learning robustness across or with respect to major stain styles, remain largely missing. Here, we present the StainStyleSampler, a toolkit for (i) the exploration and modeling of stain style variations, and (ii) the automated sampling of images or styles capturing the core components of this variation. The tool enables the extraction of various color features and deconvolved stain components, visualization of such features directly or after dimensionality reduction, modeling of style distributions using binning, clustering, and density mapping, and automated sampling of the most representative reference images. We believe that this software will equip pathologists and computer-scientists with a more versatile set of tools that can aid substantially in both the exploration and sampling of stain variation across whole slide images.

## 1. Introduction

In computational pathology, stain variation represents one of the most formidable obstacles towards the development of robust and generalizable machine learning (ML) algorithms, since model performance often drops substantially on images with appearances not covered by the training dataset [1, 2].

Two commonly used methods for overcoming this issue are stain normalization [3, 4] and stain augmentation [1, 5, 6]. Stain augmentation is frequently used during ML training, simulating variations to increase model invariance [1, 5], while stain normalization reduces variations by transferring all images to a common color style, e.g. during ML training/inference [1], clustering [7, 8], or non-ML based image evaluations [9, 10].

To determine the need for and performance of normalization/augmentation techniques, a first step is commonly the inspection of color distributions, e.g. in the RGB [11– 13], HSI [1, 14, 15], or LAB color spaces [6], or individual stain components [12]. The quantification and visualization of the stain style or color appearance across images then allows (i) to investigate inter- and intra-batch stain variations [1, 11–17], (ii) to evaluate the effect of normalization on reducing such variation [1, 6], (iii) to probe the stain variation during image augmentation [1, 6, 18], or (iv) to detect out-of-distribution images [18].

However, while the computation and visualization of stain styles represents a cornerstone in the evaluation of stain normalization and stain augmentation, comparatively few efforts have been made at producing accessible methods for sampling the underlying stain style distribution.

To address this current gap, we present the StainStyleSampler, a Python-based toolbox for the automated sampling of references from a collection of biopsy images using a variety of strategies, including different stain style representations and different sampling methods such as uniform, density-based, and clustering-based sampling. We believe that the StainStyleSampler will prove useful to researchers and pathologists for the streamlined visualization and sampling of stain variation across datasets of digitized biopsy images.

## 2. Material and Methods

### 2.1 Approach

Figure 1 displays the overall workflow implemented in the StainStyleSampler. Starting from a collection of image patches (Fig. 1A), the method first computes a corresponding stain style representation for each patch (Fig. 1B). Subsequently, the distribution of stain styles across the entire collection of patches is investigated through e.g. density mapping or clustering (Fig. 1C), to allow a sampling of distinct color appearances (Fig. 1D), resulting in the selection of patches across the observed stain variation (Fig. 1E). The next two sections will first introduce the data utilized for developing and testing the StainStyleSampler, and then outline the individual functionalities and their implementation in the StainStyleSampler.

**Fig. 1.**
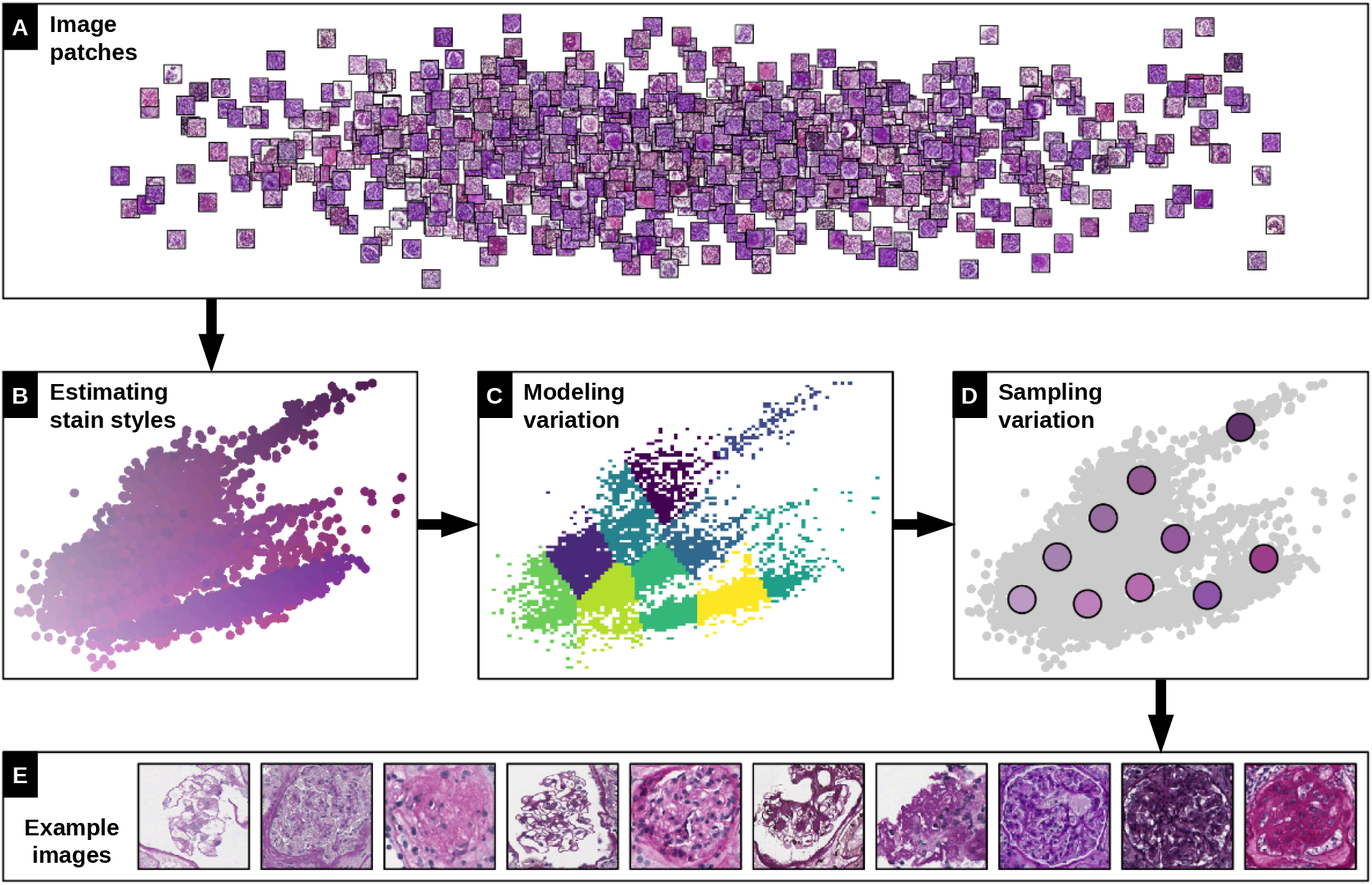
Overall pipeline implemented in the StainStyleSampler. The software takes as input a set of biopsy image patches (A), which are then first mapped to a respective stain-style representation (B). Subsequently, the distribution of stain styles is modeled or subdivided to identify different groups of appearances (C). By sampling these groups (D), it is possible to select a set of image patches capturing the overall variation in the dataset (E). The figure is based on 11054 glomeruli extracted from four publicly available kidney biopsy image repositories [19–22].

### 2.2 Data

To enable testing on a dataset with pronounced stain variation, the current study extracted patches from a broad collection of kidney whole slide images (WSIs) as described below.

#### Whole slide images

WSIs were collected from five different repositories, i.e. Bueno et al. [19, 23] (n=31), the Kidney Precision Medicine Project (KPMP, n=220)[20], the Human BioMolecular Atlas Program (HuBMAP, n=15)[21], the Kidney Pathology Image Segmentation (KPIs) challenge (n=38) [24], and the MONKEY challenge (n=80)[22]^1^.

#### Image patch extraction

From each WSI, ten patches measuring 1024 × 1024 pixels were randomly selected using HistomicsTK’s [25] *simple mask* function, with the condition that each patch contained at least 75% tissue. The selected patches then underwent a pre-processing step that involved analyzing the lower average RGB values to detect potential artifacts, such as black borders. Based on this analysis, patches exhibiting black borders were manually excluded to ensure the integrity of the dataset.

### 2.3 Implementation of the StainStyleSampler

#### Color features

To allow a flexible visualization and exploration of stain variation, the StainStyleSampler facilitates the computation of a variety of features across different color spaces, i.e. RGB (Fig. 2A), HSI (Fig. 2B) and LAB (Fig. 2C). For each color mode, the tool computes tailored statistical features: the mean and standard deviation for each channel (RGB, HSI, LAB), and skewness and kurtosis for each channel (RGB, HSI). This process can also include an optional masking of any background pixels prior to the main color analysis. Finally, these features can also be obtained after stain-deconvolution as color components of the portion of the image positive for that stain (Fig. 2D), determined using Otsu’s thresholding. While the above features are utilized for downstream feature embedding (Fig. 2E-H), the tool also always outputs the mean RGB vector for each patch, regardless of the chosen color mode, used for coloring points in embedding maps (Fig. 2I-L).

**Fig. 2.**
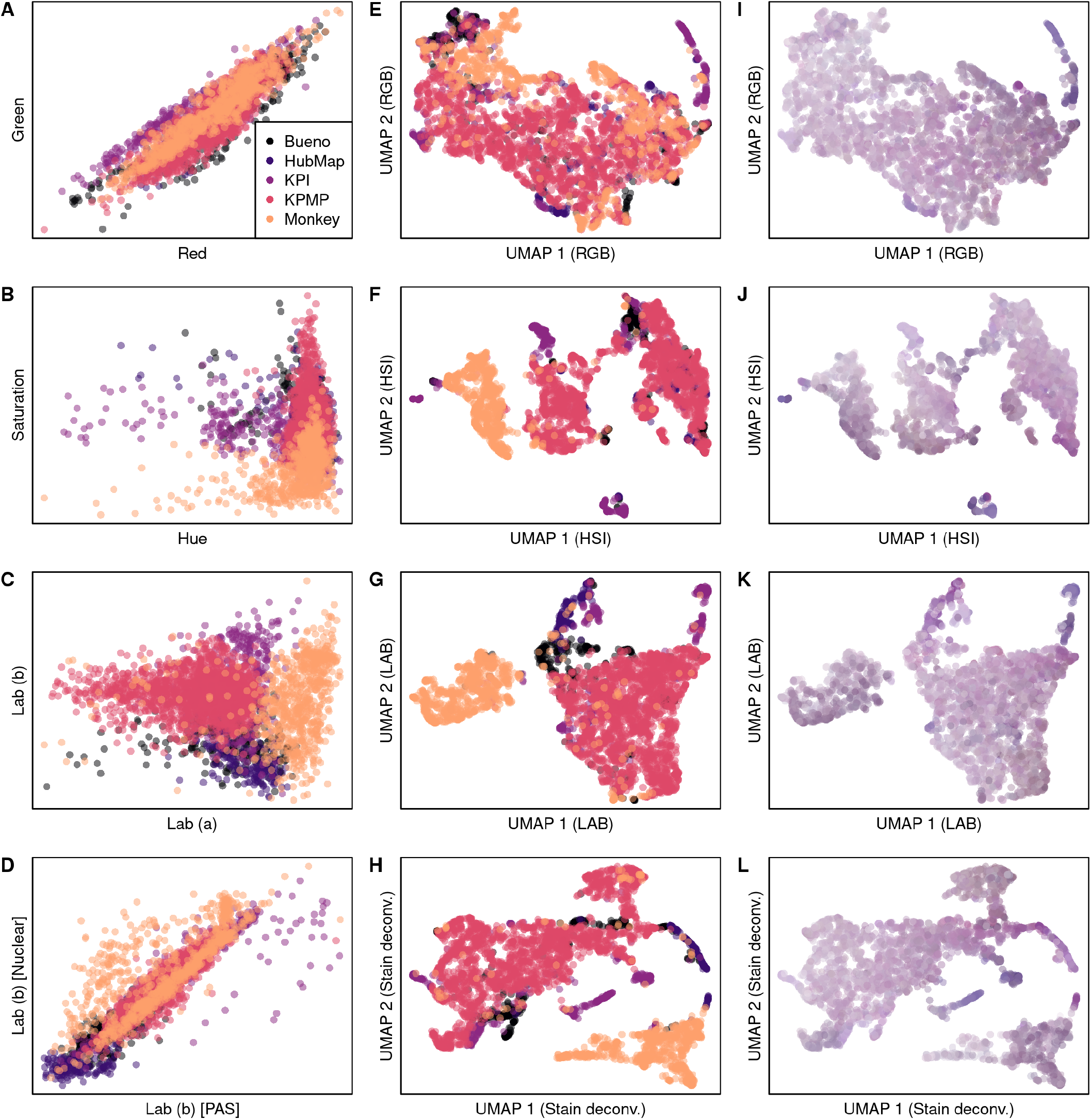
Scatter plots capturing different aspects of stain variation in the data. **A-D)** Illustration based on relationship between components from the RGB (A), HSI (B), and LAB (C-D) spaces, where (D) further integrates stain deconvolution to inspect LAB values only in pixels positive for the particular stain. Points are colored based on the originating dataset. **E-H)** Illustration of variability across datasets after UMAP dimensionality reduction of features extracted from the RGB (E), HSI (F), or LAB (G) space, or from the LAB space after stain deconvolution (H). **I-L)** The UMAP embeddings from (E-H) but colored based on the mean color per patch, illustrating the variability of colors across image patches.

#### Modeling of stain variation and Sampling strategies

The stain variation modeling and visualization process is divided into two main phases: (a) construction of embedding maps, and (b) modeling and sampling strategies.

The default embedding map utilized in the tool is obtained through a 2-dimensional Uniform Manifold Approximation and Projection (UMAP) (Fig. 2E-H), including a Principal Component Analysis (PCA) during initial data preprocessing. Alternatively, the PCA output itself may also serve as the final embedding map. To convert the continuous embedding of stain styles into a set of representative points for downstream analyses, a binned histogram is generated, and the center coordinates of the non-empty bins are extracted.

In terms of modeling strategies, four distinct strategies are implemented, grouped into three main categories, referred to as (i) Representative Targets, (ii) Density-Based modeling, and (iii) Grouped Targets, along with an additional random target approach. Representative Targets generates individual target styles using a uniform binning (Fig. 3A,B) or through clustering methods such as K-means [26] (Fig. 3C,D) or Gaussian Mixture Models [26]. The number of bins/clusters can be user-defined or automatically selected based on cluster metrics (inertia for K-means and BIC for GMM). Individual target images are then calculated as the nearest point (Euclidean distance) to the cluster centers. Density-Based modeling focuses on high-density regions within the embedding space. This is achieved by calculating contours around areas corresponding to the 90th, 95th, and 99th percentiles of the embedding maps’ histogram density (Fig. 3E). From the 99th percentile region, a specified number of images are sampled within a default distance of 1 × 10^−6^ pixels from the contour (Fig. 3F). Grouped Targets computes an average feature vector for each cluster. In this case, the afore-mentioned cluster-based methods - including K-means, GMM, and the uniform binning approach - group similar staining styles, and the target for each group is defined as an artificial template whose average and standard deviation match those of the corresponding cluster, as defined by Shen et al. [6], in order to “describe” each style. As opposed to a purely random selection of images from the dataset (Fig. 3G), the described approaches provide more control over the sampling of images with respect to the underlying distribution, e.g. to enforce the selection of reference from different areas of the stain style distribution (Fig. 3D,H).

**Fig. 3.**
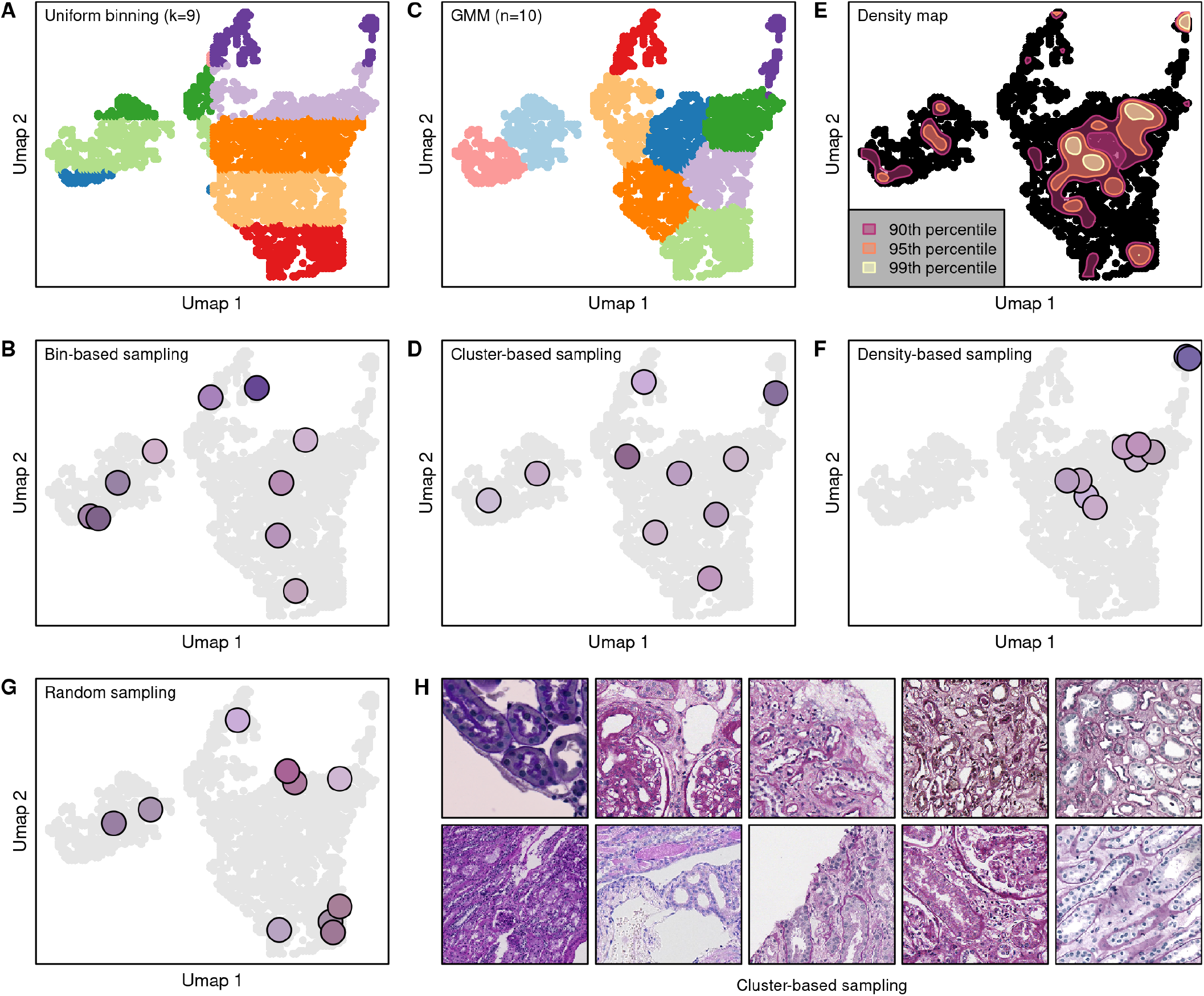
Illustration of different stain-style sampling strategies. **A**,**C**,**E)** Separating the stain style distribution into different regions using uniform bins (A), clustering (C), or density mapping (E). **B**,**D**,**F**,**G)** Selection of stain reference images based on bin centers (B), cluster centers (D), high-density regions (F), or completely random (G). **H)** Visualization of the patches selected through cluster-based sampling.

#### Libraries

The estimation, visualization, and modeling of color features proposed in this study relies in part on functions from the *HistomicsTK* [25] and *SciPy* [27] libraries for background detection, computation of statistical features, stain-deconvolution, and translation between color spaces; the Otsu’s thresholding function from the *scikit-image* library [26]; PCA and KMeans functions from *scikit-learn* [28]; and UMAP from the *umap-learn* library [29].

#### Code

The tool is publicly accessible at https://github.com/patologiivest/StainStyleSampler.

## 3. Results

When applied to a collection of kidney biopsy images from five different sources [19–22, 24], all stained with Periodic Acid Schiff (PAS), the StainStyleSampler could be used to visualize stain style variations between datasets (Fig. 2A-H) as well as illustrate the overall spread of color appearances across patches (Fig. 2I-L). Furthermore, the tool was then able to sample the underlying spread of stain styles in several ways in order to select representative image patches (Fig. 3), where the cluster-based approach appeared most effective at capturing images across all the major stain-style components in the data (Fig. 3H).

## 4. Discussion

Biopsy tissues undergo multiple laboratory processing steps - such as sectioning, staining, and scanning - before they are available as digital images for computational pathology workflows. Even minute variations during these processes can lead to substantial differences in image appearance, including changes in color, contrast, illumination, and overall quality [30]. Such variations, often referred to as stain variations, contribute to substantial batch effects, not only across different laboratories [13, 31], but also as day-to-day fluctuations within a single laboratory [32], and they usually present challenges for the training and adoption of computer-aided diagnostic tools.

Traditional strategies in stain normalization typically focus on identifying a single, most representative image either randomly and/or via an expert’s opinion as a target reference [3, 33–35]. A recent study also proposes a method to automatically select either a single global reference image, representative of the entire dataset, or class-specific references derived from class-labeled datasets [36]. However, such approaches may not capture the full spectrum of stain appearances, particularly when evaluation the robustness of ML-algorithms with respect to stain appearance, as shown in our previous work [37].

The present paper introduces the StainStyleSampler, a novel toolkit designed to automatically visualize, explore, and systematically sample reference images, aiming to capture the complete range of color variations within WSI data sets by supporting unsupervised sampling of patches from the underlying color variations via different selection algorithms, and integrating multiple visualization techniques from the existing literature into one toolkit.

## 5. Conclusion

The StainStyleSampler represents a novel tool for a versatile and streamlined visualization and sampling of stain variation. We believe that this tool might prove useful to scientists when exploring various aspects of stain-/color-related differences among histological slides and when seeking to select reference images capturing certain aspects of these differences.

## Acknowledgments

The results are in part based on data from (i) the Kidney Precision Medicine Project (https://www.kpmp.org) and (ii) the HuBMAP Program (https://hubmapconsortium.org). The project was funded by The Western Norway Health Authority (strategic research fund F-12563).

## Author contributions

*Conceptualization*: H.W.; *Data curation*: H.W., M.M.B.S.; *Formal analysis*: H.W., M.M.B.S.; *Funding acquisition*: S.L.; *Investigation*: H.W., M.M.B.S.; *Methodology*: H.W., M.M.B.S.; *Project administration*: H.W.; *Resources*: S.L.; *Software*: H.W., M.B.S; *Supervision*: H.W., S.L.; *Validation*: H.W., M.M.B.S.; *Visualization*: H.W.; *Writing – original draft*: H.W., M.M.B.S.; *Writing – review and editing*: H.W., M.M.B.S., S.L.

## Footnotes

1 Downloaded from https://registry.opendata.aws/monkey on October 21, 2024

